# Autonomous Retrieval for Continuous Learning in Associative Memory Networks

**DOI:** 10.1101/2025.05.15.654207

**Authors:** Saighi Paul, Rozenberg Marcelo

## Abstract

The brain’s faculty to assimilate and retain information, continually updating its memory while limiting the loss of valuable past knowledge, remains largely a mystery. We address this challenge related to continuous learning in the context of associative memory networks, where the sequential storage of correlated patterns typically requires non-local learning rules or external memory systems. Our work demonstrates how incorporating biologically-inspired inhibitory plasticity enables networks to autonomously explore their attractor landscape. The algorithm presented here allows for the autonomous retrieval of stored patterns, enabling the progressive incorporation of correlated memories. This mechanism is reminiscent of memory consolidation during sleep-like states in biological systems. The resulting framework provides insights into how neural circuits might maintain memories through purely local interactions, and takes a step forward towards a more biologically plausible mechanism for continuous learning.

**Author summary:** Catastrophic forgetting - when acquiring new knowledge seriously degrades previously learned information - remains a fundamental challenge in machine learning, affecting systems from simple associative networks to complex language models. One widely studied approach to mitigate forgetting is memory rehearsal, in which prior stored information are periodically replayed, an idea supported by neurophysiological evidence of memory consolidation during sleep. In this work, we show that networks with plastic inhibitory connections can spontaneously recall stored memories without external guidance. This built-in retrieval mechanism allows for the incorporation of new memories while preserving older ones, opening a path toward a biologically inspired solution to the problem of catastrophic forgetting.

## Introduction

Continuous Learning (CL) refers to a system’s ability to maintain performance across multiple tasks when operating in environments that evolve over time, requiring adaptation to changing data distributions. To do so, the learning mechanism should avoid uncontrolled forgetting of previously acquired knowledge when adapting to new information or contexts. In associative memory networks, this challenge arises when storing new activity patterns sequentially deteriorates existing memory representations, a phenomenon called catastrophic forgetting.

It is important to note that this work specifically addresses catastrophic forgetting in the context of sequential learning. This approach addresses a different challenge than the well-studied spin glass phase transitions that occur in Hopfield networks at high memory loads.

Memory rehearsal is a method that addresses the challenge of catastrophic forgetting by periodically retraining the model on previously stored patterns. This process reinforces older memory representations, preventing their degradation when new information is incorporated [1, 2]. In the mammalian nervous system, spontaneous memory replays occur during sleep [1, 3–6], suggesting a biological mechanism analogous to rehearsal techniques in artificial networks [1]. This parallel raises a fundamental question: what mechanisms enable these autonomous memory replays in biological systems?

Previous research on bioinspired neural networks demonstrates that short-term synaptic depression can facilitate spontaneous rehearsal of neural assemblies [5]. However, a significant constraint of this approach is its dependence on minimal overlap between neural assemblies. The neuronal populations in these studies share few neurons, resulting in effectively decorrelated memory representations. By contrast, classical associative memory networks like Hopfield Networks can effectively store uncorrelated memories sharing many units. Some learning algorithms even allow the storage of highly correlated patterns that share a majority of their units [7]. From a biological perspective, understanding how networks implement the rehearsal of correlated populations is crucial. Neural representations found in the cortex generally recruit extensively overlapping assemblies [8, 9]. The maintenance of overlapping assemblies is widely considered essential for cortical computation, as it supports stimulus generalization and the emergence of invariant, high-level concepts in which individual neurons participate in multiple but related representations [8–10]. This problematic is of similar importance in neuromorphic engineering contexts, as highly correlated representations are an emerging feature of deep artificial neural networks [11].

How the rehearsal of correlated memories takes place autonomously on neural substrates remains a largely unaddressed question. In this work, we focus on this issue and explore its potential application for continuous learning (CL). For the sake of bio-plausibility and potential implementation in neuromorphic substrates, we shall demand that our system exhibits the following features: 1) It stores memory states that can be highly correlated; 2) During the pattern recovery, the network does not converge toward strange attractors which would constitute false memories; 3) Plasticity rules are local, meaning that the modification of synaptic efficacy can be computed in terms of its pre- and post-synaptic neuron states; 4) It is autonomous, namely, it should retrieve all previously stored patterns from its own dynamics. The network does not have access to an external list of previously recorded memory states. For the sake of requirements (1) and (3), we shall adopt a perceptron-like algorithm, inspired by the work of Diederich & Opper (1987) [7]. On the other hand, for requirement (2) we shall use continuous Hopfield Networks (CHNs) [12]. Our work demonstrates that, kept under a certain memory load, CHNs converge exclusively to stored patterns during the retrieval. This approach avoids both, the spurious state proliferation common in Discrete Hopfield Networks (DHNs) and the shortcomings of temperature parameter fine-tuning inherent to Stochastic Hopfield Networks (SHNs) [13].

The main contribution of the present work is to introduce an algorithm to address the requirement (4). A crucial feature of our approach is the use of inhibitory recurrent plastic synapses that shrink the basin of attraction of previously visited attractor states, thus allowing for a sequential and thorough search and recovery of all previously stored correlated memory states. This recovery effectively allows the rehearsal of the stored pattern for CL purposes. The dynamic of plastic inhibitory synapses is inspired by computational neuroscience work based on actual neurophysiological data [14].

## 1 Methods

### 1.1 Continuous Hopfield Network (CHN)

In contrast with the conventional DHN model [15], where neural states are defined as binary variables, in a CHN they are continuous [12]. The dynamics of the network is defined by a set of differential equations with each neuron unit described as a leaky integration:

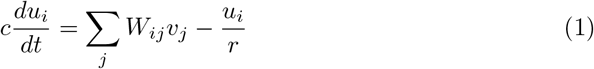

where *u*_*i*_ is the membrane potential of neuron *i, c* is the membrane capacitance, *r* is the leak resistance of each neuron, *W*_*ij*_ is the synaptic efficacy between neurons *j* and *i*, and *v*_*i*_ is the activity (or firing rate) of neuron *i* that depends solely on the potential as

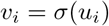

where *σ* is a monotonically increasing function of *u* with saturation to prevent runaway dynamics. We adopt 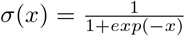. Therefore, as *σ*(0) = 0.5, each unit has a positive output at the resting state, allowing the network to have a baseline activity without external input. We shall refer to either the vector **v**(*t*) or **u**(*t*) as “states”, which should be clear from the context.

The convergence of the flow to stable states for the case of symmetric synaptic weights *W*_*ij*_ has been demonstrated [12]. Throughout this work, whenever we integrate these equations using Euler method, we do so until the network reaches convergence, defined as 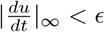, where *ϵ* = 10^*−*6^.

### 1.2 Pattern Storage and Reading

The states **v**(*t*) of the CHN evolve on [0, 1]^*N*^ through continuous dynamics, with *N* the number of neurons. Any state in this space may represent a stored pattern. For simplicity, we restrict ourselves to store binary patterns for which active neurons have a high firing rate, *v*_*i*_ ≈ 1, and inactive neurons have a low firing rate, *v*_*i*_ ≈ 0.

Hence, a pattern **x**^*µ*^ is defined as a binary vector such that 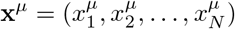 where 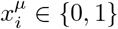 for each unit *i*. A pattern is read from the state of the network at time *t* using a threshold:

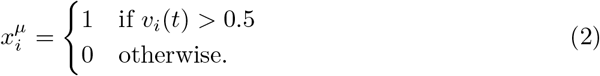

Given a binary pattern **x**^*µ*^, it is convenient to define target potentials 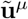 as:

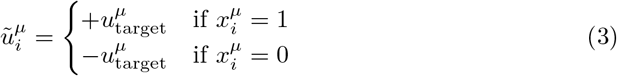

We adopt here *u*_target_ = 6 to ensure proper pattern reading following the thresholding procedure.

Inspired by previous work on DHN [7], we introduce, in Alg. 1, a perceptron-inspired learning algorithm for efficient storage of correlated patterns in CHNs. The algorithm minimizes the error between the target states and the network’s equilibrium states, ensuring that each memory becomes a stable state. Gradient descent methods typically require small step sizes to prevent the optimization process from becoming unstable and to reduce oscillations around local minima. In our implementation, we select *α* = 0.0001 as the learning rate to ensure stable convergence of weight updates. The derivation of the weight update rule can be found in the Appendix. 5.1. Although a rigorous proof of convergence for the algorithm is beyond the scope of this work, we expect that arguments demonstrating the convergence of the gradient descent algorithm (GDA) in the context of DHN could be adapted for this purpose [7]. Here, we rely on numerical evidence showing the network’s ability to successfully query and revisit stored patterns.

#### Algorithm 1 Gradient descent for the storage of correlated patterns (GDA)

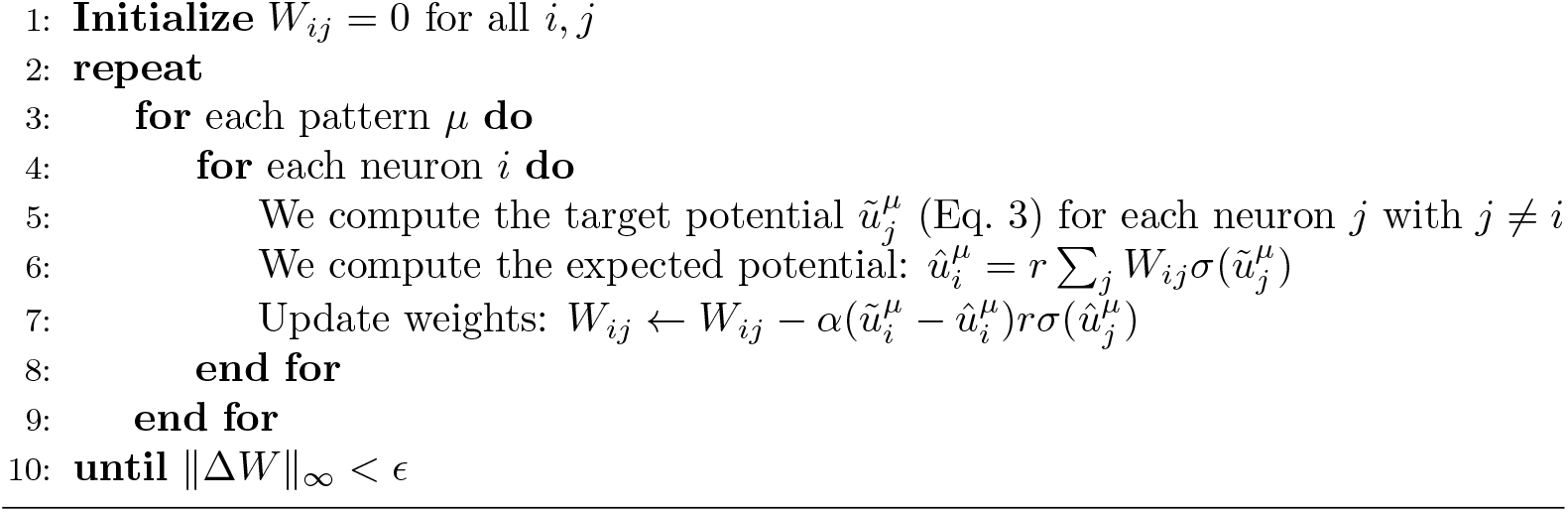

Following the network training with the GDA, each activity vector 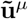 becomes an attractor. The network can now be queried as the system reliably converges to the nearest stored state from a partial cue. The corresponding binary patterns 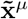 can then be accurately retrieved by thresholding at time *t*_*f*_ (Eq. 3) when the system reaches convergence. The querying procedure is detailed in Alg. 4 (Appendix 5.2) and illustrated in Fig. 1.

**Fig 1.**
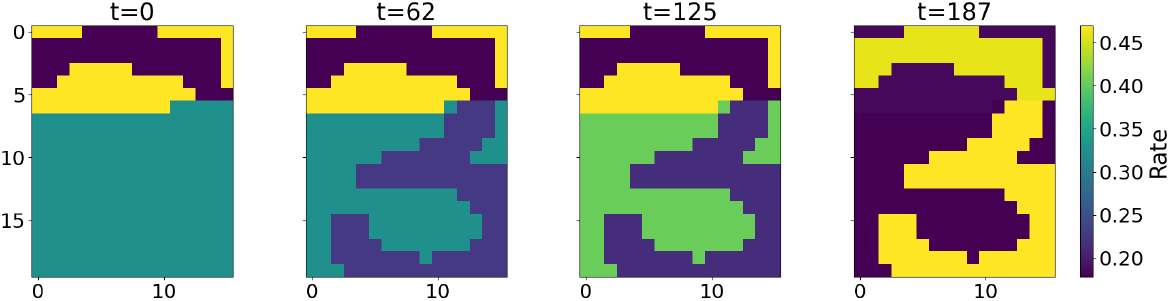
Querying of a binary-patterns representation of the handwritten digit “3” from the MNIST dataset. The network is of size 20 × 16 as each neuron codes for a pixel. From blue to yellow, the color represents the rate *v*_*i*_(*t*) of each unit. At *t* = 0 the network is initialized with the top units set to the target value as described in Appendix 5.2. The figure displays snapshots of the evolution of the rates in time for each unit of the network. The network gradually settles to the nearest stored state corresponding to the digit “3”.

It is worth emphasizing that despite the continuous nature of our model, which theoretically allows for a richer state space, we deliberately restrict ourselves to binary patterns. The interest of using a CHN is the simplicity it allows when implementing our pattern recovery mechanism (Sec. 1.4), limiting the appearance of false memories as spurious states, which are often encountered in DHNs [16]. A variant of the DHN with stochastic units, the SHN [16], would be a possible candidate to implement our algorithm, as they tend to visit only the stored patterns by properly controlling the annealing temperatures. However, the stochastic properties of these networks would require a more complex setup.

### 1.3 Continuous incorporation of correlated memories

While the GDA effectively enables the storage of correlated memories, its implementation in Alg. 1 reveals a significant limitation: it requires multiple iterations over the entire set of patterns to achieve convergence. Without the ability to reprocess all patterns, the network would suffer from catastrophic forgetting, where learning a new pattern in isolation rapidly erodes previously stored memories [1, 17–19]. By repeatedly processing all patterns, the algorithm can find a weight matrix **W** that properly separates the patterns, despite their correlations [7]. Adding a new pattern, therefore, requires access to all previously stored patterns from an external source.

This requirement for external access to the complete memory dataset stands in contrast to biological learning systems, which must incorporate new information while maintaining past memories without relying on an explicit external copy of the already stored data. To overcome this external dependency and move towards more biologically plausible learning, a solution is to develop a mechanism that allows the network to internally recover its stored memories.

The development of such an autonomous retrieval mechanism would allow us to exploit the GDA’s ability for the continuous incorporation of correlated memories. The continuous learning algorithm is formally defined in Alg. 2. The act of retraining the network on the whole set is called a rehearsal. Recovering the whole set from the network to allow rehearsal is called retrieval.

#### Algorithm 2 Continuous incorporation of correlated patterns through rehearsal of the whole memory set

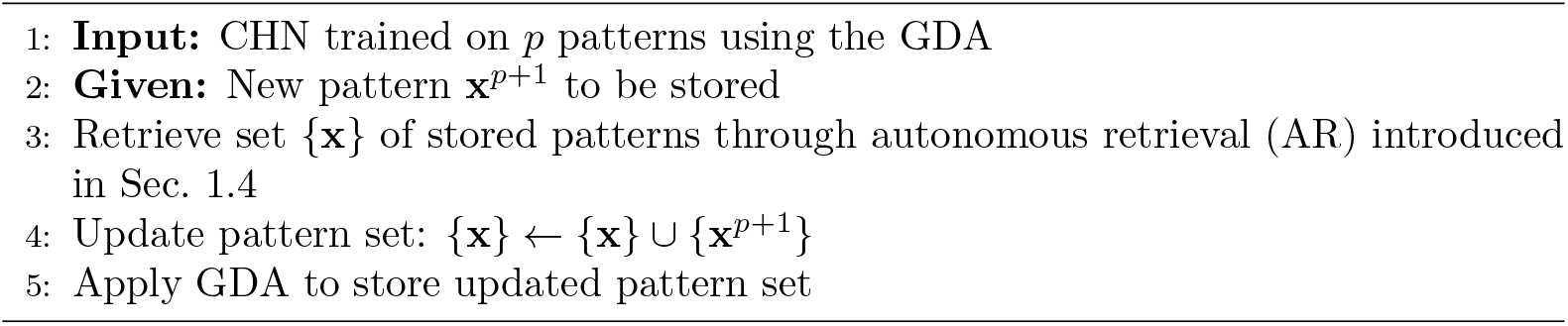

### 1.4 Autonomous retrieval

In this section, we present the autonomous retrieval (AR) mechanism allowing the recovery of stored correlated patterns in a network.

Given a trained network initialized at the “neutral” state, *u*_*i*_ = 0 for each unit *i*, the network dynamics described by Eq. 1 converge deterministically to a given stored attractor, which is thus “retrieved”. This attractor can be seen as the dominant attractor from the neutral state. The goal now is to allow for the exploration of other states to permit a complete recovery of the stored memories. To do so, we introduce a set of plastic inhibitory synapses 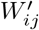.

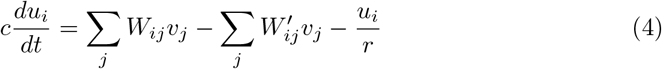

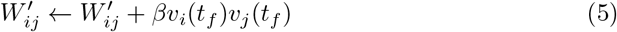

with *v*_*i*_(*t*_*f*_) the rate of neuron *i* after convergence of the dynamics. *β* represents the potentiation strength of the inhibitory synapses and is selected as a small value, typically below 0.005.

After each visited attractor, i.e. memory retrieved, its basin of attraction is made smaller by the inhibitory Hebbian-like plasticity (Eq. 5) of the synapses (*W*′). Once a pattern has been inhibited, the probability for the network to converge into it from the neutral state is reduced. By resetting the network to the neutral state after each convergence-inhibition cycle, we allow the sequential recovery of stored patterns.

Fig. 2 illustrates the sequential recovery of two stored patterns and the modification of their attractor basin during the procedure. The geometry of these attractor basins explains why a minimal inhibitory influence, resulting from a small *β* value, is sufficient to alter the trajectory. The neutral state resides near the separatrix that divides the attractor basins. Consequently, even a slight modification of the separatrix position, caused by inhibitory potentiation, can significantly redirect the network’s trajectory.

**Fig 2.**
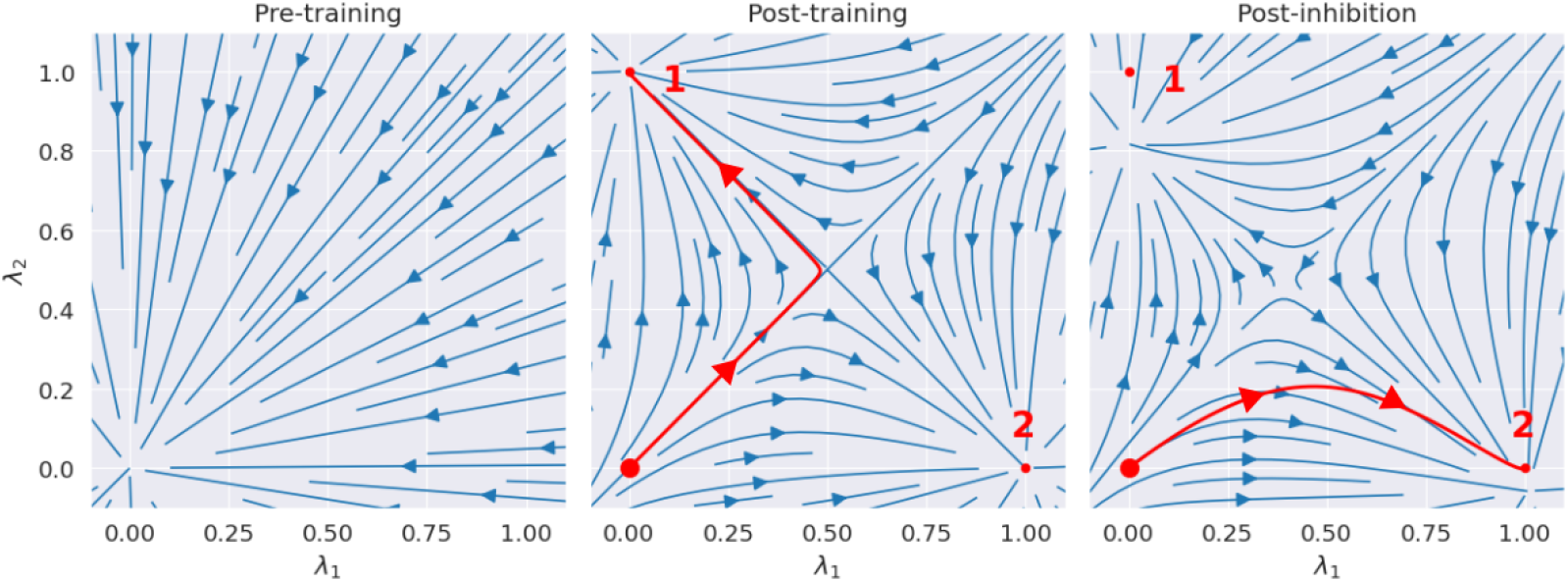
Stream plots of the CHN in a 2D subspace spanned by two pattern states. We examine a network with two stored patterns, labeled “1” and “2”, and their stable states 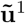 and 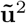 these define a two-dimensional subspace parametrized by 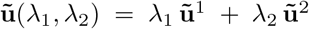. At each point 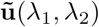 in this subspace, we compute the full *N*-dimensional time derivative via Eq. 4 and project it back onto the 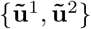-plane to obtain the plotted flow. 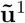 and 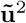 have minimal correlation as they are generated from random binary patterns **x**^1^ and ∈ {**x**^2^ 0, 1 ^*N*^}. **(left)** Before training. **(middle)** After storing the two patterns using the GDA, each pattern state (small red dots) becomes a stable attractor; the neutral state **u**_*N*_ = **0** (large red dot) is on an unstable manifold. The red line corresponds to the trajectory of the network projected on the subspace. The trajectory follows the unstable manifold before arbitrarily falling into one of the two stored patterns. **(right)** *W*′ is updated to apply enhanced inhibition to pattern 1 which shrinks its basin of attraction. This induce a shift in the position of the separatrix, favoring the flow to pattern 2. An exaggerated large inhibitory coefficient *β* = 0.3 is used to illustrate the modification of the vector field. Similar stream plots are obtained from networks storing patterns with various degrees of correlation using the GDA.

The potentiation of inhibitory synapses tends to distort the stable states associated with stored patterns. As this distortion is minimal for small *β*, it is mainly compensated for by the thresholding mechanism used to read the network output (Sect. 1.2). To further reduce the impact of this distortion, we divide the convergence dynamic into two phases, the “biased” phase and the “free” phase. The biased phase guides the network to converge toward states that have not been retrieved. It corresponds to the simulation of the network with inhibitory synapses, Eq.4, until convergence. The free phase then allows the network to complete its convergence to an undistorted stored state. It corresponds to the simulation of the network without inhibitory synapses (Eq.1) until convergence. The whole procedure is detailed in Alg. 3.

Fig. 3 provides a visualization of AR for binary-patterns representation of handwritten digits from the MNIST dataset [20]. For the first iteration, the inhibitory drive is null as no pattern has been retrieved. The pattern corresponding to the binary picture of a 3 is recovered. The network state is reinitialized with the updated inhibitory matrix (**W**′). The network now inhibits the recovery of a 3, which induces convergence toward a second pattern, here a 6. The inhibitory matrix (**W**′) is updated again and now inhibits both the 3 and the 6 together. The last pattern, corresponding to a 5, is then recovered.

**Fig 3.**
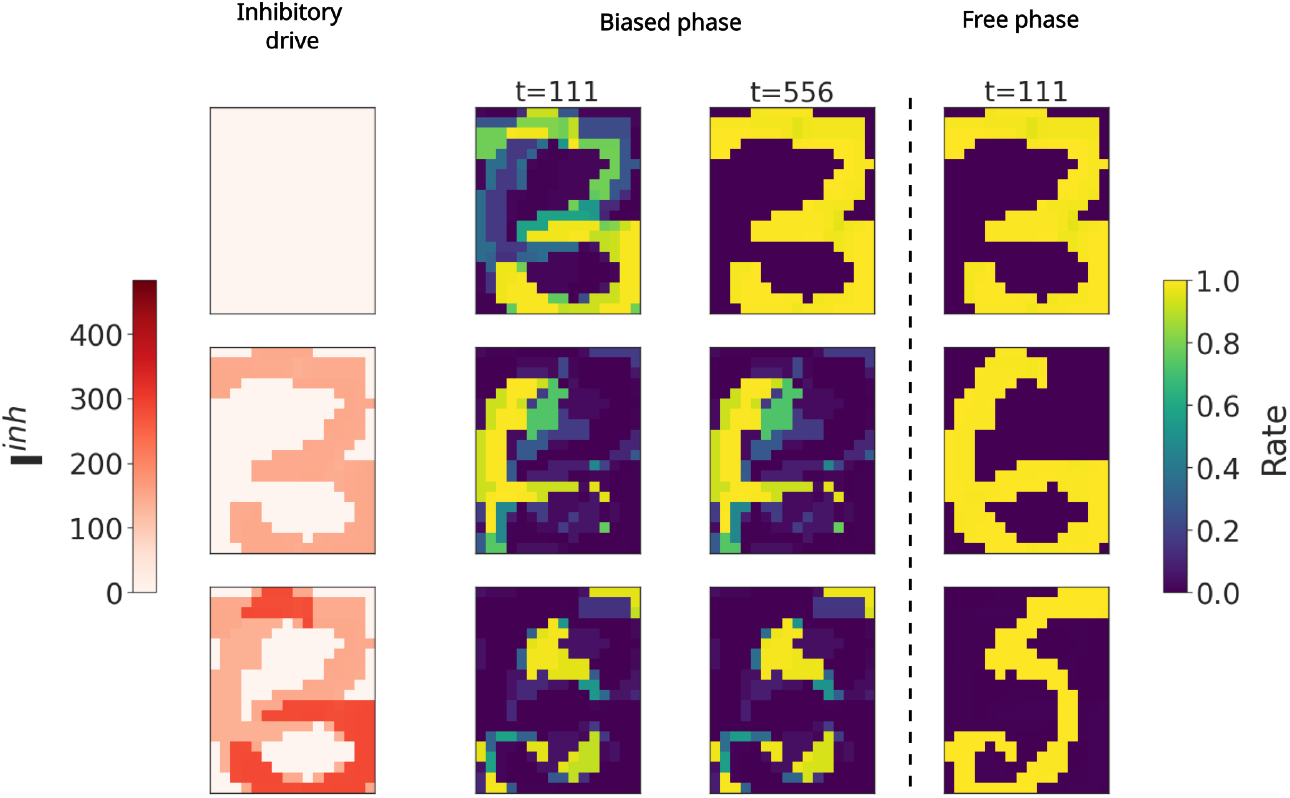
Recovery of three binary-patterns representations of handwritten digits from the MNIST dataset, “3”, “6”, “5”. The network is of size 20 × 16 as each neuron codes for a pixel. Left panels (in red): Evolution of the inhibitory drive to each neuron, defined as 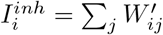. Right panels (blue and yellow): snapshots of the evolution of the rates *v*_*i*_(*t*) for each unit of the network. Each row corresponds to the start of a new memory retrieval in the CHN. The black dashed-line corresponds to the removal of inhibitory weights W’, i.e. change from biased to free phase, as described in Alg. 3. The panels illustrate how the biased phase “orients” the evolution, and then the free phase provides the precise convergence to a stored pattern. An exaggerated large inhibitory coefficient *β* = 0.25 is used to illustrate the stable states deformation at the end of the biased phase, highlighting the need of the free phase.

The order in which patterns will be visited is an emerging feature of the learning algorithm that hasn’t been studied in this work. With each iteration of the AR algorithm, inhibitory synapses undergo potentiation. This growth is constrained only by the number of iterations and the value of *β*. Theoretically, such an unbounded growth could cause the weights in **W**′ to reach magnitudes comparable to those in **W**, potentially disrupting the convergence dynamic. However, in practice, this potential issue can be managed by adjusting the value of *β* based on the number of iterations. Specifically, when more iterations are required, using a smaller *β* is sufficient to maintain robust pattern retrieval without compromising attractor dynamics.

In Fig. 4, we illustrate the dynamics of the CHN during the retrieval of stored memory patterns in networks with various loads. The load corresponds to *M/N*, with *M* the number of stored memories, and *N* the size of the network. For low loads, see Fig. 4(top), the system converges sequentially to the attractors corresponding to the stored patterns. Once every memory has been recovered, if the simulation is not ended, the network falls back into the stored states without showing any false memories. The lower the value of *β*, the longer this dynamic can proceed without encountering spurious states. The free phase has no utility in this scenario, the attractors are well separated, and the disruption of the energy landscape by the inhibitory plasticity is minimal. For critical loads, see Fig. 4(middle), the retrieval process becomes more challenging. The attractors exhibit reduced separation, and the network struggles to converge to the stored states once inhibition is applied. In this scenario, the free phase demonstrates its utility. At the end of the biased phase, during the second iteration of the AR, the network remains in a mixed state characterized by ambiguous correlations with multiple stored patterns. The free phase enables the network to resolve this ambiguity and ultimately converge to a stored state. At high loads, see Fig. 4(bottom), the network successfully retrieves some stored states, but mostly falls into spurious attractors. Under these conditions, even the free phase cannot rescue the recovery dynamics.

**Fig 4.**
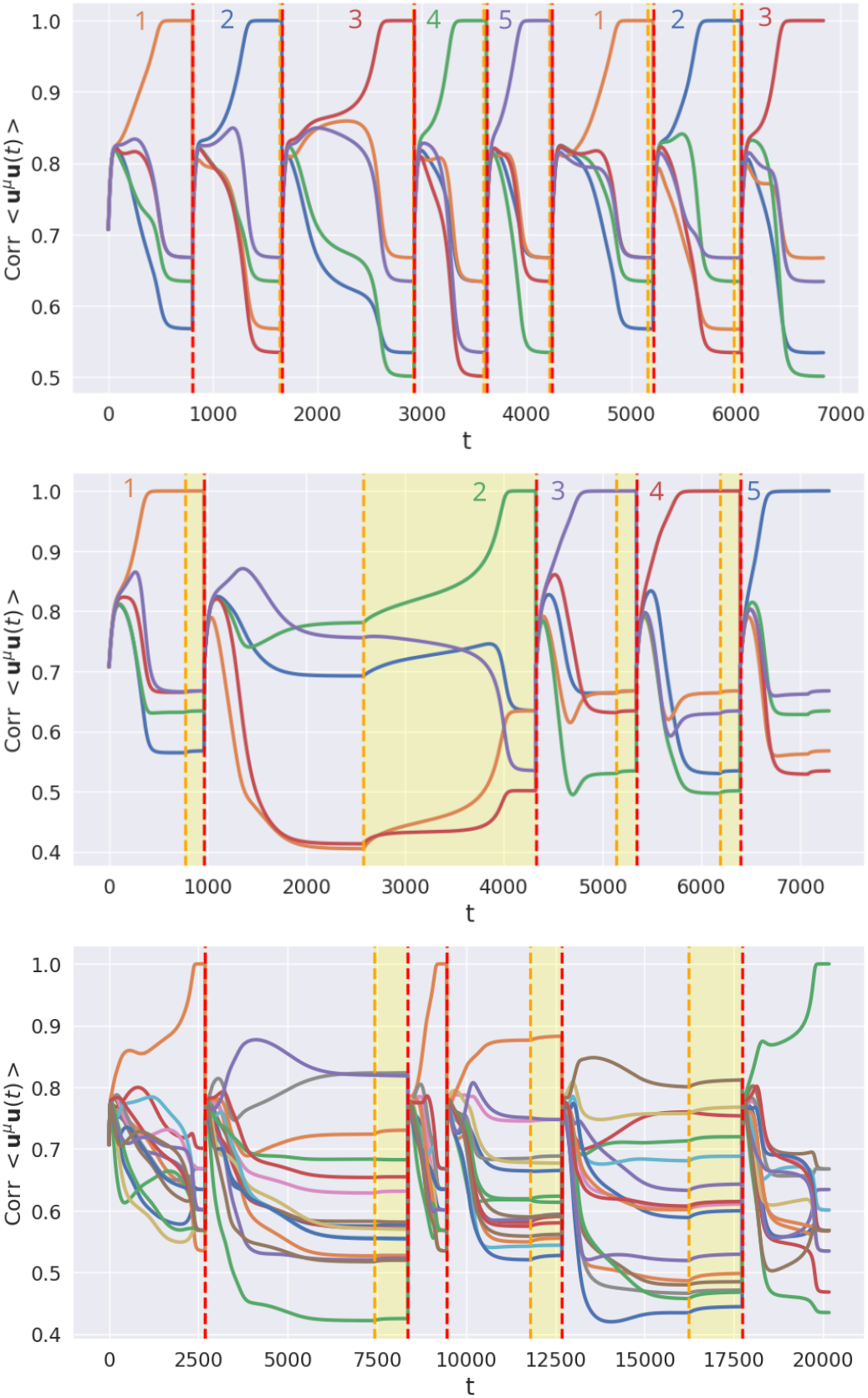
Visualization of AR (Alg. 3). The x-axis shows the number of time steps for simulations of the CHN, with all query iterations of AR concatenated. The y-axis shows the correlation between the state of the network and any stored state, with each color corresponding to a specific memory. Vertical red dashed lines separate successive memory retrieval sequences. At the beginning the matrix W’ is updated and the initial state is the neutral one. The yellow dashed lines indicate the end of the biased phase. Between yellow and red lines is the free phase, during which the W’ matrix is momentarily set zero, to converge precisely into a stored state. Considered patterns have minimal correlation as they are generated from random binary vectors. **(Top)** Recovery dynamics for a low-load network: 5 patterns for a network of 60 units. **(Middle)** Recovery dynamics for a critical-load network: 5 stored patterns for a network of size 30. **(Bottom)** Recovery dynamics for a high-load network: 16 stored patterns for a network of size 60.

These autonomous recovery dynamics are reminiscent of memory replays observed in the central nervous system of mammals during quiescent states, such as sleep or rest. This process is believed to function as a consolidation mechanism that mitigates or modulates memory forgetting [1, 3–6].

#### Algorithm 3 Autonomous retrieval (AR)

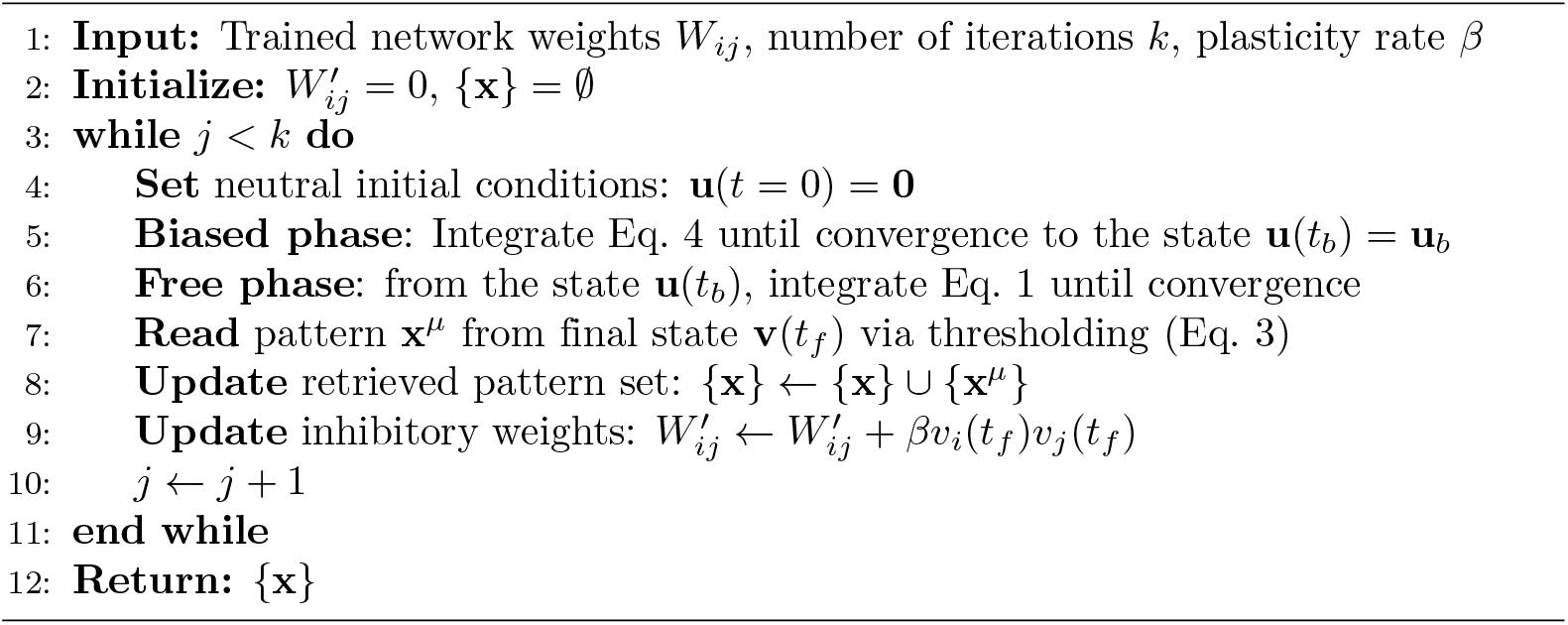

## 2 Results

In this section, we evaluate the ability of our algorithm to retrieve correlated patterns from a given network. We will consider the retrieval of pattern sets with various amounts of correlation. Each set gets assigned a random binary pattern, the parent pattern. Each pattern of a given set is generated by randomly choosing and randomizing a fraction (1 − *ρ*) of bits from the parent pattern as described in Alg. 5 (App. 5.3). Therefore, a higher *ρ* induces more correlation, while a lower *ρ* results in less correlation. Networks with various loads will be tested.

Fig. 5 illustrates, for various correlations, the loads for which AR allows systematic recovery of all stored patterns without external cues or memory lists. This property enables the continuous incorporation of new correlated memories, as described in Alg. 2. The retrieval improves with network size - larger networks can reliably retrieve more stored patterns without encountering false memories. We can observe the rather surprising feature that a moderate amount of correlation (*ρ* ≈ 0.5) tends to improve pattern retrieval compared to highly correlated and minimally correlated pattern sets. In fact, this finding contrasts with traditional associative memory models, where correlation typically degrades performance [21].

**Fig 5.**
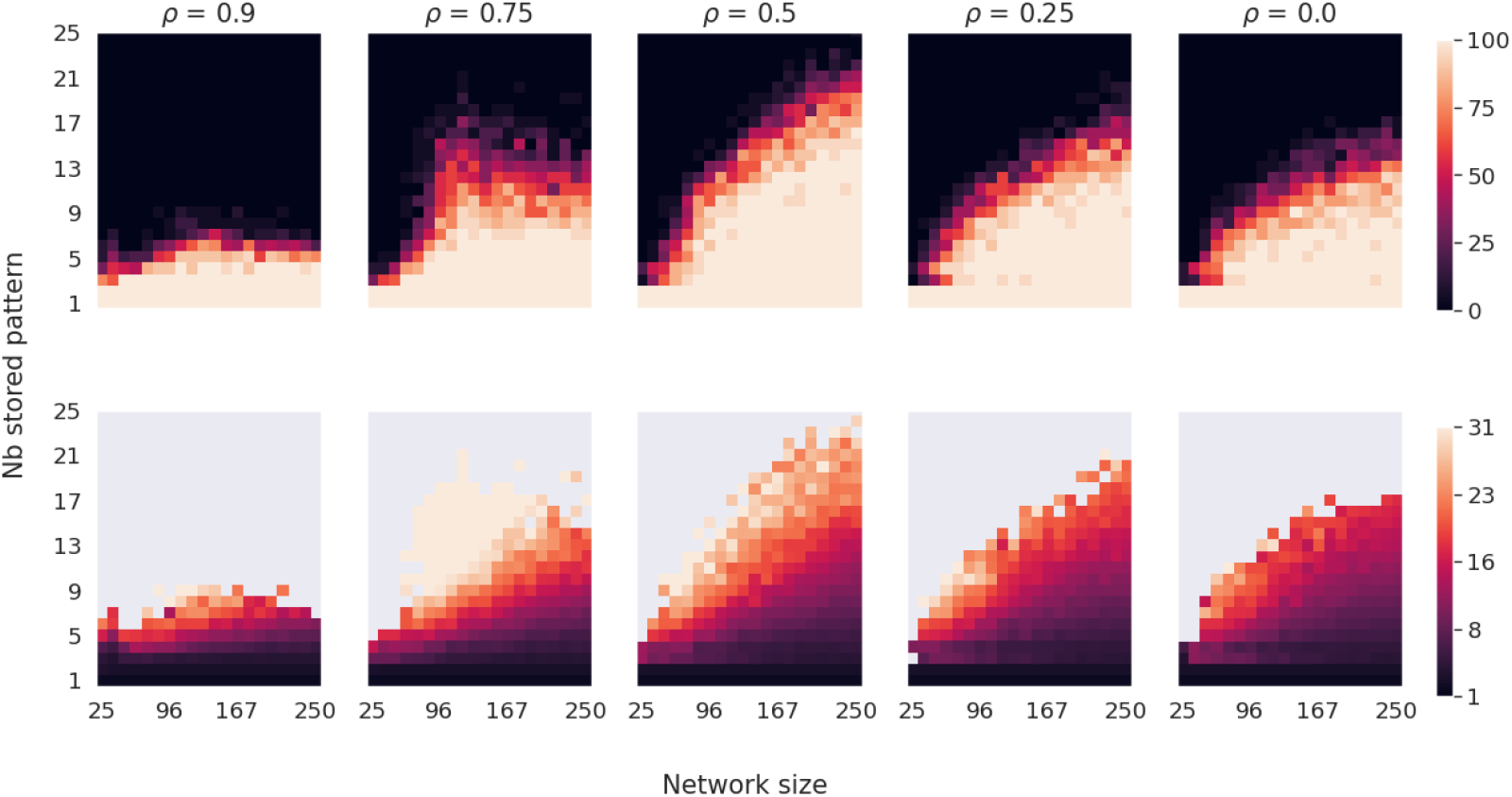
Recovery capacity of AR for various correlations, modulated by *ρ*. Data are averaged over 20 simulations with different pattern sets. Correlated patterns are generated using Alg. 5 (Appendix 5.3). For networks of various sizes and numbers of stored patterns, we employ Alg. 3 until all patterns are recovered or until a spurious state is found. **(Top row)** The percentage of ‘full retrieval,’ i.e., simulation runs where no spurious state occurs before recovering all stored patterns. **(Bottom row)** Number of iterations required to recover all patterns. Recovery is best for *ρ* = 0.5, which denotes equal number of correlated and uncorrelated bits. For small *ρ* values with few correlated bits, performance worsens. For highly correlated sets, only very low loads are recovered without false memories.

Our interpretation is that an intermediate amount of correlation helps the system to be driven in the “good” direction in early stages of the evolution, i.e. while still rather close to the neutral state, and the energy landscape is rather featureless. Once driven to a point of the state space proximal to all the stored patterns, the system can finish the convergence. As illustrated in Appendix 5.4, Fig. 8, this push towards the good direction can be measured through the average correlation between the synaptic drive of each unit and the stored patterns.

As emerges from our simulations, to limit the appearance of false memories, the values of *β* must be relatively small compared to the target potential *u*_target_. In our case, we found it convenient to keep *β* smaller than 0.005. By keeping the nudging of the convergence dynamic small, the emergence of false memories that might otherwise result from deformations in the energy landscape is reduced. Fig. 6 indicates the existence of a trade-off: Smaller *β* values require more iterations to retrieve all patterns but provide greater stability, while larger values accelerate pattern retrieval at the cost of increasing the probability of encountering a spurious state. Higher values of *β* than those considered in this study lead to catastrophic degradation of recovery dynamics for which no stored patterns are recovered. Lower values of *β* only lead to the need for more iterations when recovering the pattern set.

**Fig 6.**
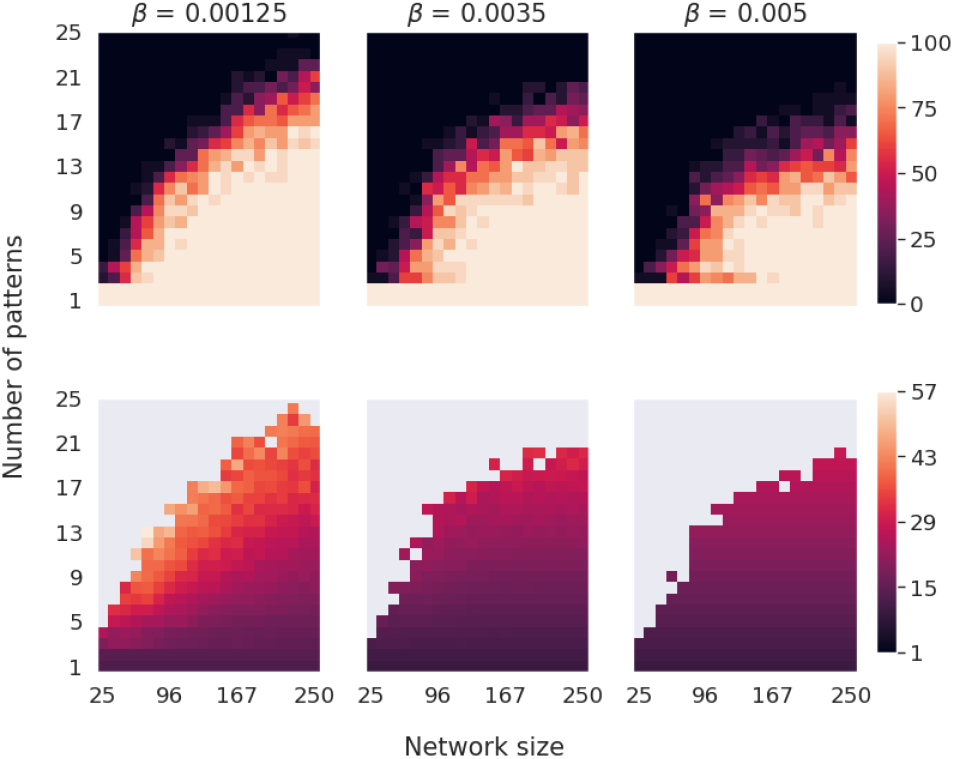
Similar to Fig. 5 but for various *β* values. Correlated patterns are generated with a *ρ* of 0.5. Lower *β* values require more iterations to recover the complete set of stored patterns. Higher *β* values diminish the number of iteration needed but increase the probability of finding spurious state when the load is important.

We shall now describe some qualitative information on the efficiency of our algorithm. As expected, the number of iterations required to successfully store patterns using Alg. 1 increases with the memory load (Fig. 7). Moreover, the higher the correlation between patterns, the higher the number of iterations needed for storage. Overall, our method requires significantly more iterations than previously documented for associative memory tasks in DHNs [7]. We can argue that this increase in computational cost may come from two factors. First, training a CHN through GDA (Alg. 1) inherently requires more iterations than training DHNs. Second, we observed that a smaller convergence parameter *ϵ* in Alg. 1, while computationally more demanding, yields superior retrieval performance. We hypothesize that this tighter convergence criterion induces stronger competition between pattern attractors at the neutral state. This competition results in enhanced network responsiveness to subtle modifications of attractor basins induced by W’ during retrieval. These observations led to the adoption of a very small *ϵ* = 10^*−*6^, which demanded more iterations when performing the GDA.

**Fig 7.**
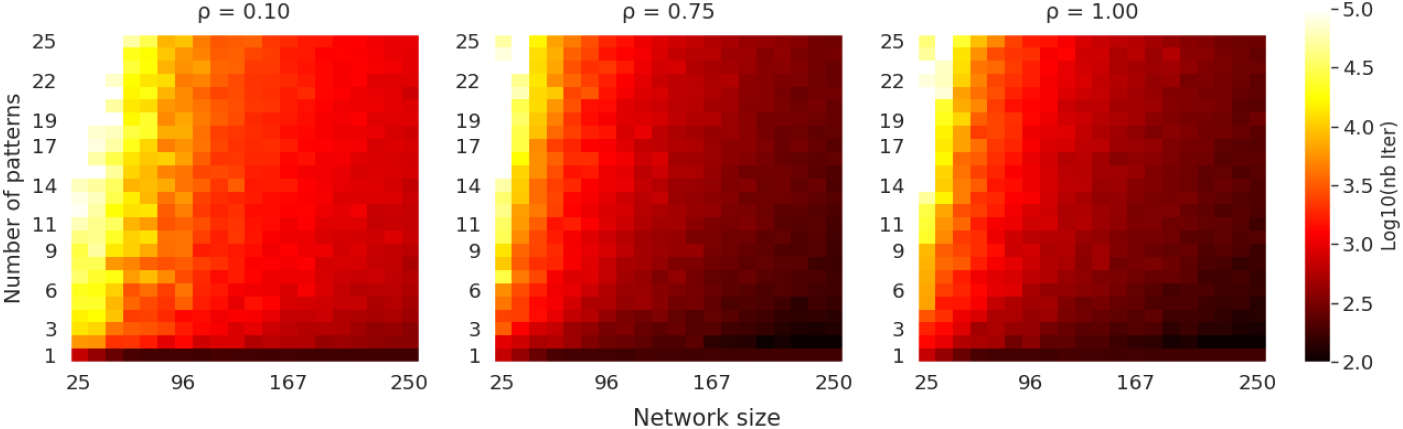
Convergence time of the GDA. Color intensity shows the number of iterations required to store a given number of patterns using GDA (Alg. 1). Data are averaged over 20 simulations for various pattern sets.

## 3 Discussion

Our work introduces a biologically inspired mechanism for the continuous incorporation of correlated patterns in associative networks. CL is made possible through the autonomous recovery of all stored patterns during a retrieval phase. By systematically retrieving memories, the network can incorporate new patterns while mitigating the forgetting of the ones already stored. Autonomous retrieval is made possible by inhibitory plastic synapses, avoiding the necessity of recalling patterns from an external list.

Previous work in computational neurosciences indicates that inhibitory circuits may play a critical role in the regulation of neural activity and plasticity [14, 22]. Here we demonstrate that inhibitory plasticity could be one of the key mechanisms that allow the sequential reactivation of memories observed during sleep and resting states [4, 23]. A property of associative networks highlighted by our approach is that subtle changes in inhibitory connectivity can drive substantial shifts in network dynamics without disrupting the fundamental structure of stored attractors. Inhibition, therefore, allows context-dependent activity of the network as observed in experimental setups [24]. Biological neural circuits might, therefore, employ similar mechanisms to navigate complex, correlated, memory spaces. Our results indicate that larger networks experience fewer spurious state visits, suggesting improved reliability with scale. However, understanding how these dynamics extend to networks of biologically relevant sizes would require further investigation.

Traditional associative memory models typically suffer from decreased capacity when storing correlated patterns [16]. However, our findings revealed the rather unexpected feature that moderate correlation levels actually improve pattern retrieval in the context of autonomous retrieval. We argue how this result may arise from the way that correlated structures influence the geometry of the attractor basins and the ensuing flow towards them. A more thorough theoretical analysis of this phenomenon could provide information on the factors that influence the robustness of recovery dynamics in both artificial and biological systems.

## 4 Conclusion

In this work, we demonstrate how inhibitory synaptic plasticity can enable autonomous exploration of attractor landscapes in continuous Hopfield networks. Our key finding reveals that, under a critical load, inhibitory plasticity allows networks to systematically retrieve the entire set of stored memories, even for highly correlated sets. This property allowed us to propose and test an algorithmic scheme for continuous learning leveraging the ability of gradient descent to store correlated patterns. The capacity for self-directed pattern exploration, emerging from inhibitory modulation, offers insights for both biological memory consolidation and neuromorphic computing.

## 5 Appendix

### 5.1 Derivation of the gradient descent learning rule

At steady state, when 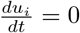, from Eq. 1 we have:

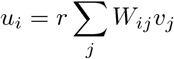

where *v*_*j*_ = *σ*(*u*_*j*_). We can then define

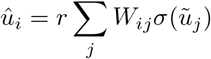

the expected potential of the unit *i* when the rest of the units are at their target potential 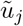.

We can then define the error function *E* as the sum of squared differences between the expected potentials 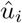 and target potentials 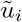.

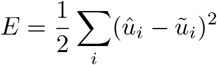

Taking the derivative with respect to the expected potential:

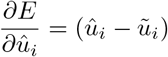

To compute how the error changes with respect to the weights *W*_*ij*_, we use the chain rule:

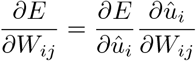

The partial derivative with respect to *W*_*ij*_ is:

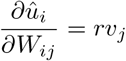

This is an approximation that captures the immediate direct effect of *W*_*ij*_ on 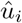, assuming that *v*_*j*_ remains constant and ignoring feedback effects in the recurrent network. This approximation is valid when the optimization steps are small relative to the nonlinearities in the sigmoid function.

Therefore, the error gradient with respect to *W*_*ij*_ becomes:

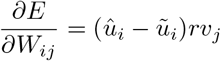

Finally, we update the weights using gradient descent:

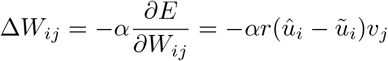

where *α* is the learning rate, typically set to 0.001 to ensure convergence.

### 5.2 Querying procedure

The network’s ability to retrieve stored patterns can be tested through partial cues. Given a subset of informed units from a pattern *µ*, we initialize their potentials according to the partial pattern while leaving other units at rest :

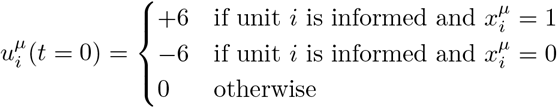

The network then evolves according to Eq. 1 until it reaches a stable state 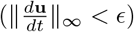. The thresholding procedure then allows for the reading of a stored binary pattern **x**^*µ*^:

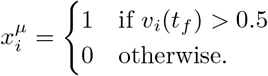

with *v*_*i*_(*t*_*f*_) the rate of neuron *i* after convergence of the dynamic. Alg. 4 summarize the procedure.

When operating below capacity, the dynamics typically converges to the stored pattern most similar to the initial cue. The final state is interpreted as a binary pattern using the thresholding procedure (Eq.3). As with traditional DHNs, retrieval success depends both on the network load and on the number of units informed in the initial cue. Although a theoretical analysis of the storage capacity under Alg. 1 would be of interest, our focus here is on the autonomous retrieval mechanism, which operates in a regime well below this limit for which pattern retrieval is highly reliable.

#### Algorithm 4 Querying with convergence of the network using Euler method

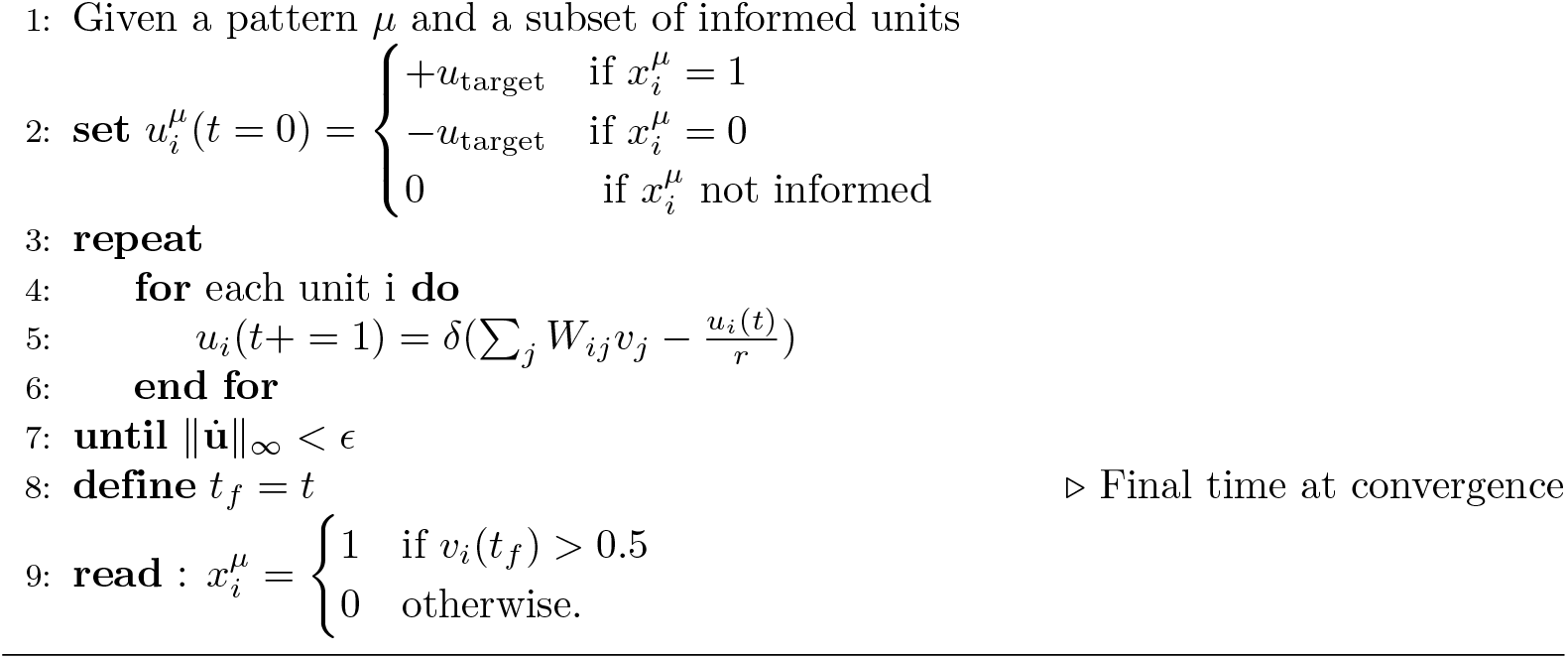

### 5.3 Construction of correlated patterns

#### Algorithm 5 Generation of *p* correlated random patterns

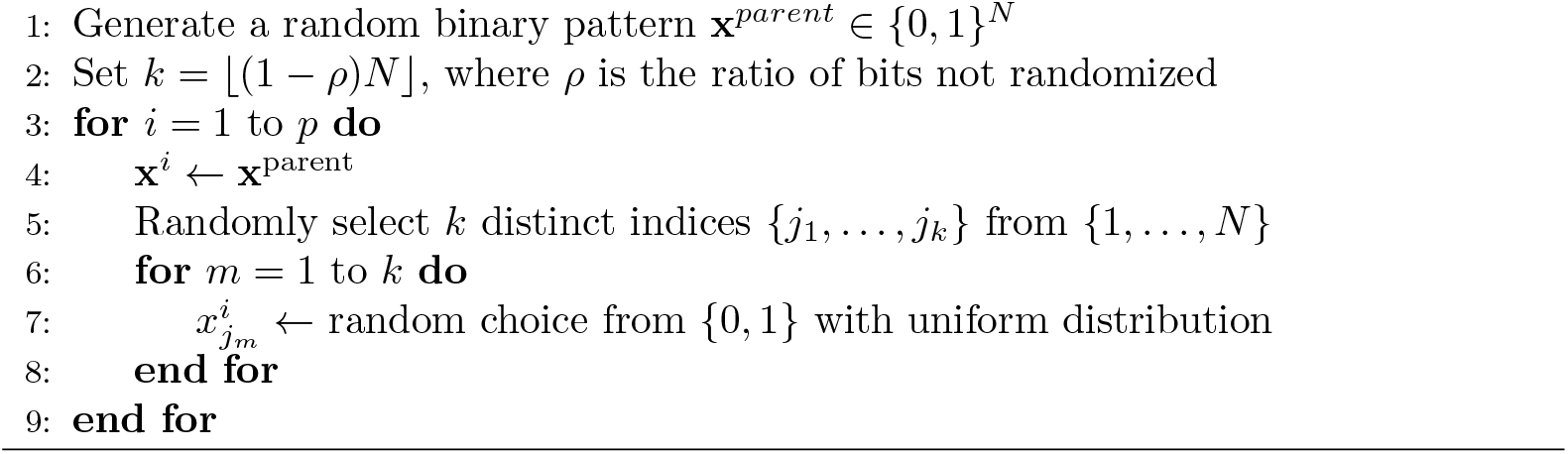

### 5.4 Drive-pattern correlation and spurious states

**Fig 8.**
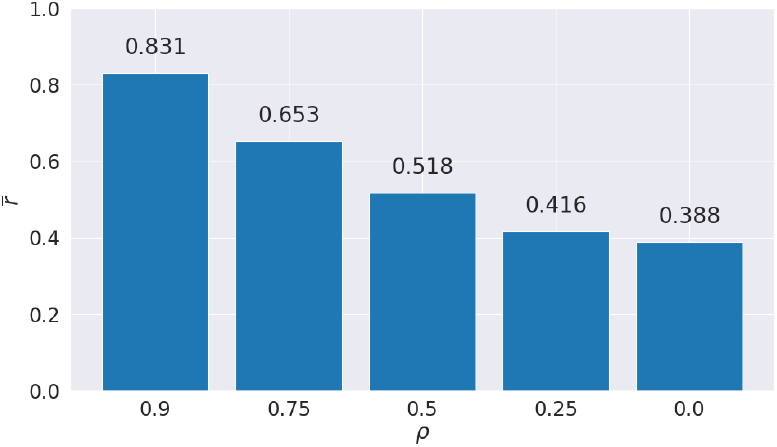
For networks trained with sets of varying correlation *ρ*, we compute the synaptic drive of each unit 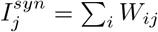. This synaptic drive is the main component influencing the dynamic of the network at the neutral state, when the leak is null. Then, we compute the average correlation between each stored pattern and the synaptic drive: 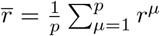 with *r*^*µ*^ = Corr(**x**^*µ*^, **I**^*syn*^). We can see that, the higher the *ρ*, the stronger the average correlation between the drive **I**^*syn*^ and stored patterns. At the neutral state, from which the network is initialized, stronger correlations induce a stronger push of the network state toward stored patterns. This could explain why our algorithm performs better with moderately correlated patterns.

### 5.5 Parameters for simulations

**Table 1.** CHN parameters.

| Name | Value | Description |
| --- | --- | --- |
| $\epsilon_{sim}$ | $10^{-6}$ | Convergence constant |
| $r$ | 1 | Leak time constant |
| $c$ | 1 | Integration time constant |
| $\delta$ | 0.001 | Euler step size |

**Table 2.**
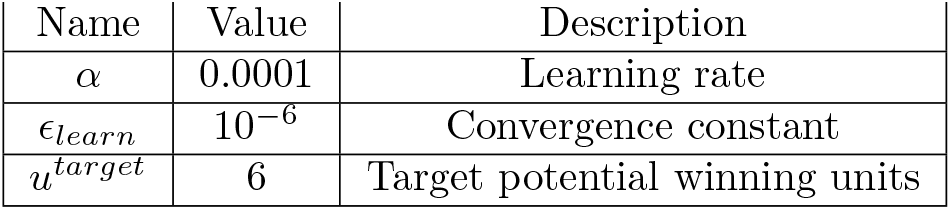
Gradient descent.

| Name | Value | Description |
| --- | --- | --- |
| $\alpha$ | 0.0001 | Learning rate |
| $\epsilon_{learn}$ | $10^{-6}$ | Convergence constant |
| $u^{target}$ | 6 | Target potential winning units |

## 6 Acknowledgments

This project has received financial support from the French DIM AI4IDF and the ANR “MemAI” project ANR-23-CE30-0040. This work was supported by Paris Region.

